# Precision 1070 nm Ultrafast Laser-induced Photothrombosis (PLP) of Depth-targeted Vessels *in vivo*

**DOI:** 10.1101/2022.06.22.497150

**Authors:** Liang Zhu, Mengqi Wang, Peng Fu, Yin Liu, Hequn Zhang, Anna Wang Roe, Wang Xi

## Abstract

The cerebrovasculature plays an essential role in neurovascular and homeostatic functions in health and disease conditions. Many efforts have been made for developing vascular thrombosis methods to study vascular dysfunction *in vivo*, while technical challenges remain, such as accuracy and depth-selectivity to target single vessel in the cerebral cortex. Herein, we first demonstrates the evaluation and quantification of the feasibility and effects of Rose Bengal (RB)-induced photothrombosis with 720-1070 nm ultrafast lasers in raster scan. A flexible and reproducible approach is then proposed to employ a 1070 nm ultrafast laser with spiral scan for producing RB-induced occlusion, which is described as precision ultrafast laser-induced photothrombosis (PLP). Combine with two-photon microscopy imaging, this PLP displays highly precise and fast occlusion induction of various vessel types, sizes, and depths, which enhances the precision and power of the photothrombosis protocol. Overall, the PLP method provides a real-time, practical, precise, and depth-selected single-vessel photothrombosis technology in cerebral cortex with commercially available optical equipment, which is crucial for exploring brain vascular function with high spatial temporal resolution in brain.

## Introduction

The cerebral vasculature forms an intricate and highly interconnected vascular network of arteries, capillaries, and veins, allowing blood to flow through and meet the high metabolic demands of the brain^1-4^. Although similar vascular network characteristics can be found throughout the cortex, there are variations across the cortical layers and between different cortical areas^5,6^. Understanding this difference in vascular topology and hemodynamics goes hand in hand with regional spatiotemporal neuron activation^7,8^, which is the basis of a multitude of biomedical imaging techniques such as functional MRI^9,10^ and intrinsic optical imaging^11,12^. In addition, dysfunction of the cerebrovasculature and insufficient blood supply will generate irreparable damage to the brain tissue and induce cerebrovascular diseases and neurodegenerative disorders^1,5,13-16^. There has been a long-standing interest in the fine topological architecture and hemodynamics regulation of vascular blood flow in health and disease states. However, the degree of the precise spatial micro-scale of vascular signals correlate with local neural activity has been difficult to assess^17,18^. There are many efforts devoted to developing the precision *in vivo* approach for vascular regulation, mainly by occluding the vessels, including surgical suturing or ligation, such as middle cerebral artery occlusion (MCAO)^19-21^, and thrombosis induced by fluorescent microspheres^22^, endothelin-1^19^, even magnetic nanoparticles^23^. Although these methods have constructed various vascular occlusion models, technical challenges remain in terms of depth, accuracy, and reliability. Additionally, there are few efficient thrombosis methods for the single vessel, especially parenchymal capillaries, due to the small size and elusiveness^24^.

Light-induced vascular photothrombosis is one of the precisely controlled occlusion methods^25-27^. Single-photon illumination is the most commonly used method to produce occlusions by activating the intravascular Rose Bengal (RB), owing to its reproducibility and flexible selectivity. RB has been widely utilized in vascular photothrombosis research, as it could rapidly generate reactive oxygen species (ROS) with the excitation of a green laser, causing platelet aggregation and eventually occluding the vessel^25,27-29^. However, when exposing the underlying vasculature to light, the out-of-focus excitation could lead to undesired blockage of adjacent vessels, especially the overlying vessels, due to the nature of the one-photon reaction. The main constraint of one-photon-based excitation approaches remains the limited spatial resolution and penetration depth: short-wavelength and wide-field illumination make it hard to target an individual capillary at the deep layer^30^. Recent studies have proposed solutions for RB-excitation in parenchymal capillaries. They employed longer-wavelength ultrafast lasers (720, 1000nm) to photoactivate RB in the cerebral vasculature, producing shallow-layer parenchymal single-vessel photothrombosis^31,32^. An alternative method has also been proposed to induce parenchymal occlusions by photothermic damage of endothelial cells using infrared ultrafast laser pulses^25,26^. Despite these investigations, a reliable protocol for precision single-vessel occlusion in the deep layer, as its efficiency, accuracy, and penetration depth could be affected by the ultrafast laser’s adjustable parameters is needed.

In this study, we evaluated and quantified the efficiency of RB-induced coagulation cascade with ultrafast laser at different wavelengths from 720 nm to 1070 nm combined with two-photon laser scanning microscopy (TPLSM). TPLSM is an important tool to study the dynamics of blood flow and metabolism *in vivo*, due to its superior depth-penetration in the tissue and focus-specific fluorescence excitation^33,34^, which enables long-term observations with high spatiotemporal resolution *in vivo*. We proposed a novel and highly reproducible approach of precision ultrafast laser-induced photothrombosis (PLP) of single vessels with RB using a 1070 nm laser. This repeated PLP paradigm employed spiral scan for RB excitation, allowing fast, precise and controlled occlusion of the target single vessel while monitoring the related vascualature networks in real-time. Our approach enhances the power of single-vessel occlusion paradigm, providing high success rates of occlusion in various vascular types (capillary, artery and vein), sizes (3-500 μm diameter), and depths (up to 800 μm), even in awake mice.

## Results

### Compare the efficacy of target photothrombosis induced by raster scan of ultrafast lasers at different wavelengths

To evaluate the photothrombosis efficacy at different wavelengths of ultrafast laser at different wavelength, we first imaged vasculatures using TPLSM system paired with a tunable titanium-sapphire laser. The vasculature was labeled with fluorescein-dextran (2 MDa FITC) to verify the structure and hemodynamic changes during photothrombosis processing. Our paradigm includes 20 min of baseline vasculature imaging, followed by an up to 5-minute photothrombosis period and then a post-thrombosis imaging period (Fig. 1c). During baseline imaging, the target vessel was chosen and imaged with the excitation of a 920 nm laser. For photothrombosis, the excitation laser was modulated at different wavelengths (720, 780, 860, 920, 1000, 1070nm) and then coupled into the microscope system (Fig. 1a). The excitation beam was focused into the target vessel through a cranial window using a water immersion objective (Fig. 1b). Considering the difference in tissue penetration characteristics of lasers at different wavelengths, all attempts were carried out with the same power (80mW) and targeted the capillaries that lie under the cortical depth of 80-100 μm and range from 3 to 8□µm in diameter. Once the target vessel was selected, a dose of ∼50 μL RB solution was injected into the plasma, and then raster scan was performed up to 5 mins since the fast clearance from circulation of RB^35^. The ROI for the target vessel segment was manually selected and quickly refocused with the excitation laser. During photothrombosis processing, a repeated raster scan was delivered and restricted in the ROI while imaged in real-time at 40 to 80 fps simultaneously. Scanning was terminated when clots formed and the flow stopped over 10 s.

**Figure 1.**
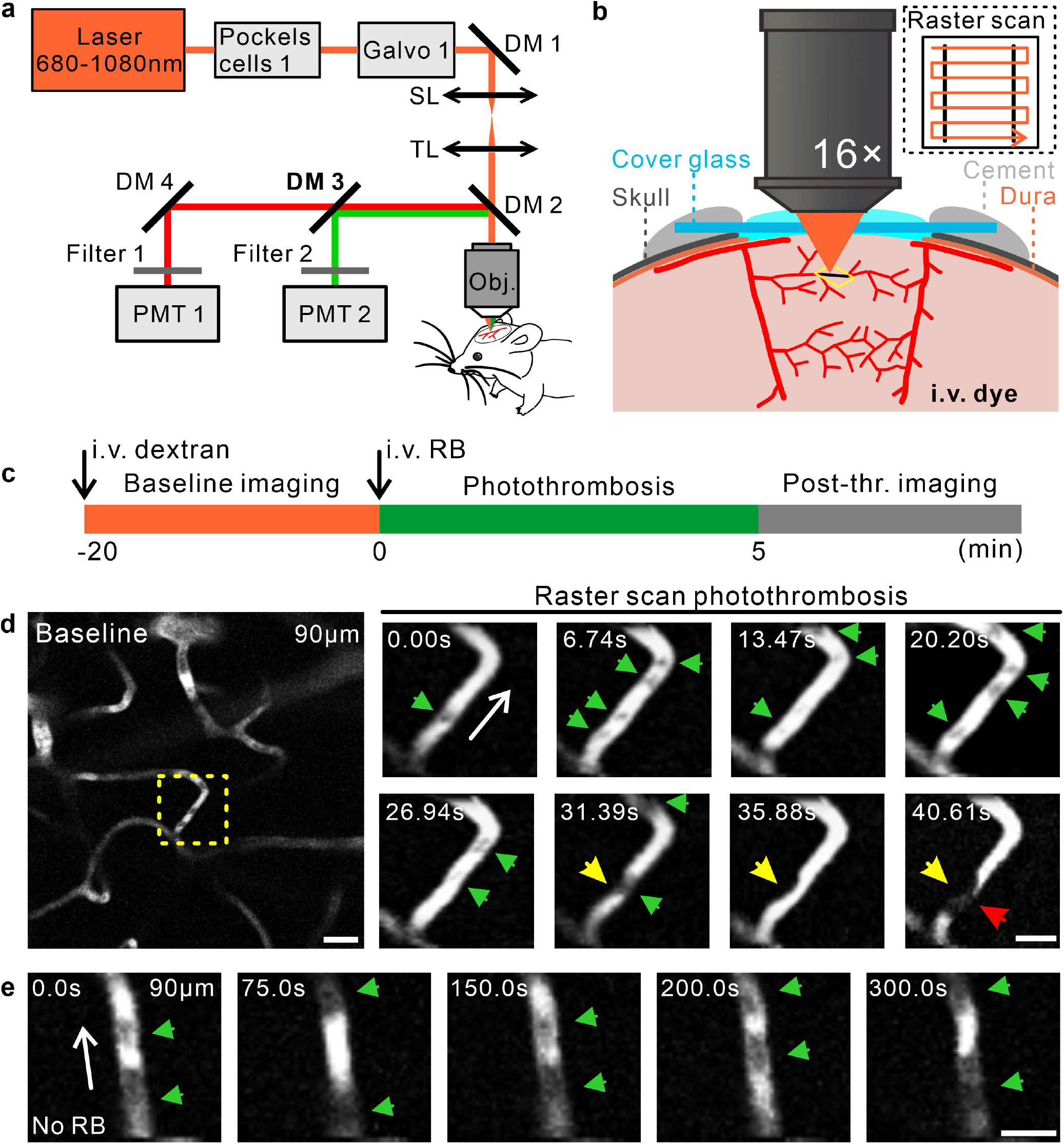
Illustration the RB-induced photothrombosis with 720 nm-1070 nm ultrafast laser in raster scan model *in vivo*. a. Schematic of the photothrombosis configuration with TPLSM. The optical path for the excitation laser (orange) was sketched. Galvo 1: galvometers; DM 1-2: dichroic Mirror; SL: scan lens; TL: tube lens; PMT 1-2: photo-multiplication tubes; Obj: objective. b. Schematic, laser was focused by the objective through a cranial window and controlled as raster scans in the target ROI including the target vessels labeled with RB. c. An experimental time course for photothrombosis of targeted vasculature. d. A photothrombosis example of a parenchymal capillary at 90 μm depth. Images on the right show snapshots of real-time planar imaging during raster scan photothrombosis, enlarged from the yellow rectangle in the baseline image. The green arrows indicated the red blood cells which passed through the vessels’ lumen as black plaques. The yellow arrows indicated the platelets aggregation and attachment to the vessel wall, with a clot (red arrow) formed at 40.61 s. Scale bar: left image: 20 μm; right images: 10 μm. e. A control vessel at 90 μm depth without RB during 300s raster scans. The green arrows indicated the red blood cells which passed through the vessels’ lumen as black plaques. The white arrow corresponds to the flow direction. Scale bar: 10 μm.

Our example showed a phothrombosis process of a capillary at 90 μm depth (Fig. 1d). The target vessel segment received a repeated raster scan of a 720 nm laser and was imaged in real-time. Initially, red blood cells passed rapidly through the vessel’s lumen, appearing as black plaques in the image (green arrows). Thereafter, platelets were attached to the vessel wall, leading to a “focal contraction” of the vessel segments (yellow arrows). Although the RBC has great deformability in facilitating the passage through this “constriction”^36^, it eventually got stuck at the clot site and induced an occlusion (red arrow) as the clot enlarged (Supplementary Movie 1). To verify whether this photothrombosis was induced by sorely energy deposition effects of the ultrafast laser, we applied the same scan pattern to the no RB-labelled vessels in same depth. The control vessel displayed neither attachment of platelets nor blood cells aggregation in the vessel’s lumen during the 300 s’ scan (Fig. 1e, Supplementary Movie 2).

We next tested the performance of the photothrombosis effects at different wavelengths of ultrafast lasers in two cortical layers. We examined 137 subjected capillaries at 80-100 μm from 16 mice with laser power at 80 mW. All the wavelengths of lasers were able to reliably induce occlusion in RB-labelled vessels in the majority of cases, as each group shows a high success rate above 78% (Fig. 2a) (Success rate, 720 nm: 84%; 780 nm: 78.26%; 860 nm: 90.48%; 920 nm: 90.48%; 1000 nm: 86.96%; 1070 nm: 87.5%). Notably, during the raster scan of photothrombosis, BBB leakage occurred in a few capillaries only with excitation of shorter wavelength (Fig. 2a) (BBB leakage rate, 720 nm: 8%; 780 nm: 13.04%; 860 nm-1070 nm: 0%). We then defined the period from the start of the scan to the time of complete cessation of blood flow as occlusion time to quantify the photothrombosis effeteness at different wavelength lasers. Notably, the occlusion formation costs significant longer periods with the excitation lasers at 860 nm and 920 nm (860 nm: 106.3 ± 41.97 s; 920 nm: 123.3 ± 42.49 s; Mean ± SD). Other four wavelengths of lasers (720, 780, 1000, 1070 nm) exhibited a more rapid thrombotic process (Fig. 2c) (720 nm: 48.10 ± 21.09 s; 780 nm: 66.09 ± 38.05 s; 1000 nm: 55.25 ± 35.32 s; 1070 nm: 65.01 ± 38.79 s; Mean ± SD).

**Figure 2.**
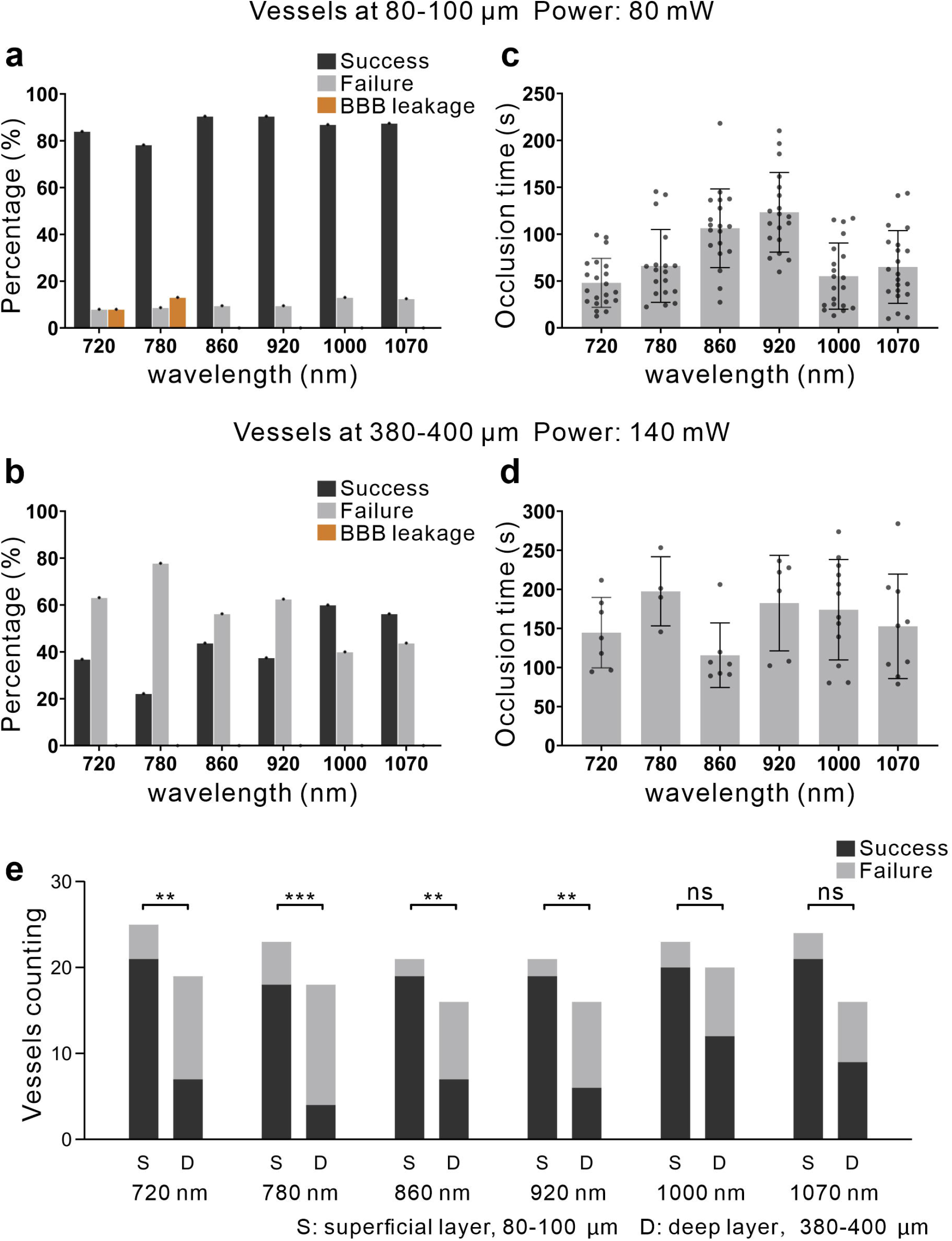
Quantification of the success rate and occlusion time of photothrombosis at different laser wavelengths. a, b. Column plot of percentage of three categories (success, failure, BBB leakage) capillaries after up-to-300 s raster scan at different wavelengths in two cortical layers (a: 80-100 μm, N= 137 vessels from 16 mice; b: 380-400 μm, N= 105 vessels from 12 mice). c, d. Column plot of occlusion time at different wavelengths in two cortical layers. Each data dot represented the occlusion time of an individual capillary. (c: 80-100 μm, N= 118 vessels from 16 mice, Mean± SD; d: 380-400 μm, N= 45 vessels from 12 mice, Mean± SD.) e. Comparation of capillary occlusion success/failure ratio in two cortical layer. Grouped in per each wavelength. (p= 0.0018 for 720 nm; p= 0.0005 for 780 nm; p= 0.0034 for 860 nm; p= 0.0011 for 920 nm; p= 0.0781 for 1000 nm; p= 0.0588 for 1070 nm; Fisher’s Exact Test)

Considering the distinct tissue penetration properties of laser at different wavelengths, we explored the photothrombosis effects at deeper cortical layer. We examined 118 subjected capillaries at 380-400 μm from 12 mice with increased laser power at 140 mW. Relative to the superficial layer, all groups showed worse photothrombosis effects as groups with shorter wavelength showed a lower success rate around 40% while 1000 nm and 1070 nm group greater reliability (Fig. 2b) (Success rate, 720 nm: 36.84%; 780 nm: 22.22%; 860 nm: 43.75%; 920 nm: 37.5%; 1000 nm: 60%; 1070 nm: 56.25%). Interestingly, there was no sign of BBB leakage occurred in all groups during the raster scan of photothrombosis (Fig. 2b) (BBB leakage rate, 720 nm-1070 nm: 0%). Notably, the occlusion time between each group showed no significant difference in success samples (Fig. 2d) (720 nm: 144.6 ± 45.09 s, n = 7; 780 nm: 197.5 ± 44.26 s, n = 4; 860 nm: 115.7 ± 41.34 s, n = 7; 920 nm: 182.4 ± 61.18 s, n = 6; 1000 nm: 174.0 ± 64.23 s, n = 12; 1070 nm: 152.7 ± 66.88 s, n = 9; Mean ± SD). In addition, we compared the success/failure attempts in superficial and deep layers with each laser to characterize the occlusion effects with penetration properties (Fig. 2e). The results showed that the 720, 780, 860, and 920 nm lasers had significantly lower success/failure probability in these two layers, suggesting that the energy lose due to penetrating the cortex while applying raster scan in deep cortex (p= 0.0018 for 720 nm; p= 0.0005 for 780 nm; p= 0.0034 for 860 nm; p= 0.0011 for 920 nm). The 1000 and 1070 nm lasers showed no significant difference between these two layers, suggesting that the limited energy loss and the power could still support for efficient photothrombosis in the deep cortex (p= 0.0781 for 1000 nm; p= 0.0588 for 1070 nm).

### Precision and depth-selective photothrombosis by spiral scan of 1070 nm laser

Repeated raster scan in target ROI can effectively produce RB-induced parenchymal photothrombosis, however, the thermal damage of the vessel walls caused by constant scan and, high laser power demand for deep cortical occlusion give cause for concern. Recent work has demonstrated that different from raster scan, spiral scan enables more efficient heat dissipation^30^, which helps to prevent the photothermic damage while ensuring fast and efficient excitation. In addition, previous works have indicated the excellent tissue penetration properties of 1070 nm lasers^37-39^, allowing the minimum attenuation in the brain tissue, which reduces out-of-focus excitation with lower focused power. We proposed an improved method (PLP) of spiral scan photothrombosis that employed a 920 nm laser for imaging and a 1070 nm laser for RB excitation. The imaging laser was emitted by a tunable titanium-sapphire laser and then coupled into the microscope system (Fig. 3a, orange path). The 1070 nm laser was emitted by a non-tunable femtosecond laser and modulated to spiral scan by another galvanometer (Fig. 3a, green path with spiral). During the baseline imaging period, the target vessel was chosen and mapped at appropriate magnification with the imaging laser, and then a circle ROI was drawn in the vessel’s lumen, whose size was matched to the target vessel while avoiding covering the vessel wall. While applying spiral scan, the 1070 nm laser scanned at a constant ultra-fast speed from the center outward to the ROI edge repeatedly with specified revolutions throughout the duration (Fig. 3b). The number of revolutions was fixed to 0.5 μm per circle, and the laser power was controlled from 80 mW to 200 mW under the objective altering to the specific situation. To efficiently photoactivate RB without thermal damage caused by constant scan, we separated imaging and photoexcitation procedures, and formulated a photothrombosis paradigm of 3 seconds scanning duration followed by 2 seconds of imaging repeatedly up to 5 min (Fig. 3c).

**Figure 3.**
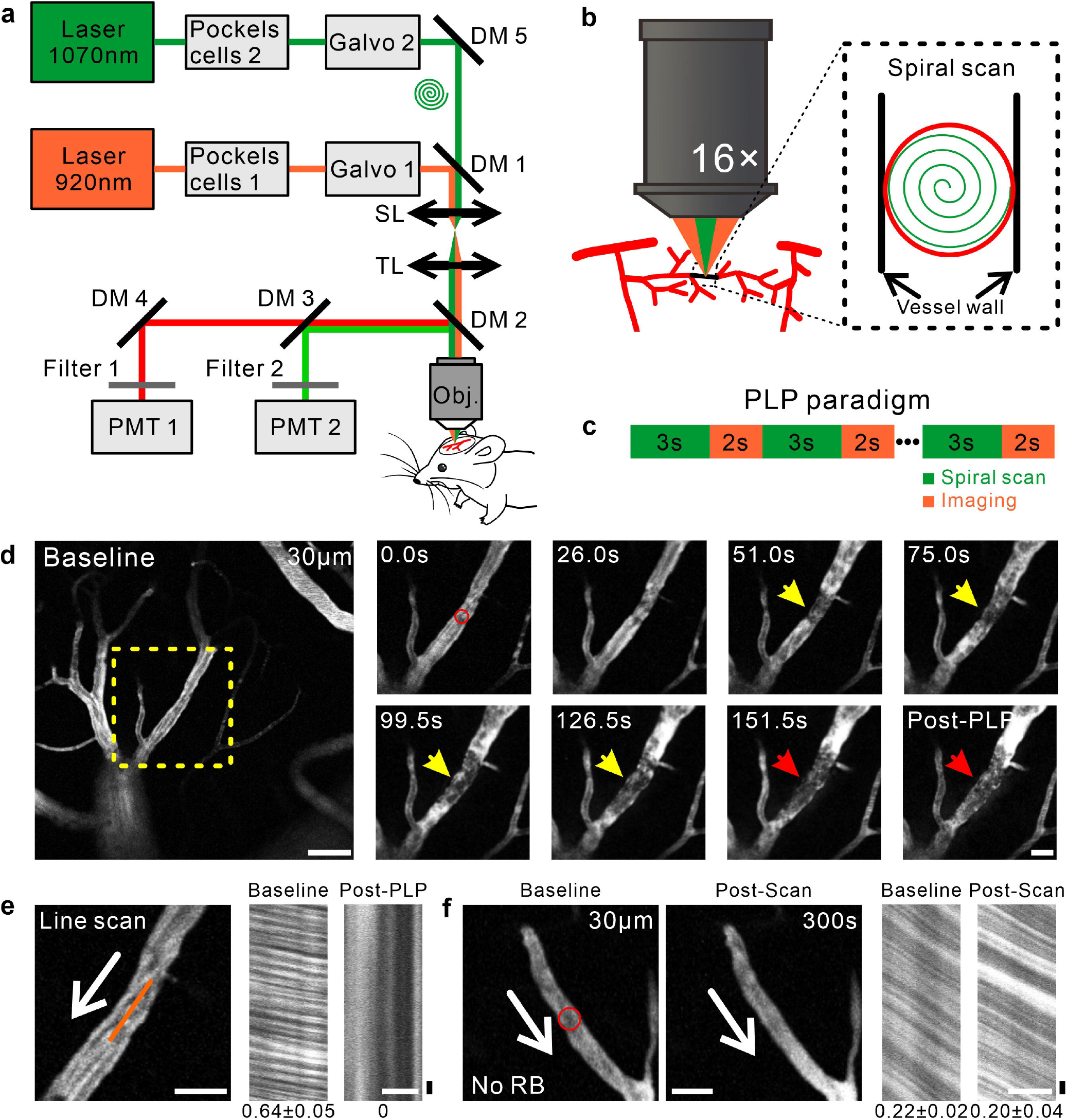
*In vivo* RB-induced photothrombosis with spiral scan at 1070 nm (PLP). a. Schematic of the spiral-scan photothrombosis configuration with TPLSM. The optical paths for two-photon imaging (920 nm, orange) and ultrafast photothrombosis (1070 nm, green) were sketched. The two scan patterns were modulated by galvanometers separately. Galvo 1-2: galvometers; DM 1-5: dichroic Mirror; SL: scan lens; TL: tube lens; PMT 1, PMT 2: photo-multiplication tubes; Obj: objective. b. Schematic, 1070 nm laser (green) was focused by the objective and modulated as spiral scans in the target ROI (red) inside the vessel’s lumen. c. The PLP paradigm with repeated 3s spiral scan following 2s imaging was sketched. d. A photothrombosis example of a pial vein. Images on the right showed snapshots of real-time planar imaging during clot formation, enlarged from the yellow rectangle in the baseline image (left image). The red circle in the first slice displayed the target ROI to deliver spiral scans. The yellow arrows indicated the formation of a clot, with a clot (red arrow) formed at 151.5 s. Scale bar: left image: 50 μm; right images: 20 μm. e. Line-scan image of the target vessel of baseline and post-thrombosis. Left: The orange line indicated the scanned line in target vessel. Right: The line scan image and data in baseline and after PLP. Unit, mm s^-1^; horizontal scale bar, 20 μm; vertical scale bar, 0.01s. f. A control vessel at 90 μm depth without RB after spiral scan. The red circle in the first slice displayed the target ROI to deliver spiral scans. In planar e and f, the white arrow corresponded to the flow direction. The number below each line-scan image indicated the flow velocity. Unit, mm s^-1^; horizontal scale bar, 20 μm; vertical scale bar, 0.01s.

The example showed a PLP process of a pial vein (Fig. 3d, Supplementary Movie 3). The pial vein was chosen as the target vessel and then the ROI was drawn in the vessel’s lumen (red circle, diameter = 12.1 μm). We then injected RB solution into the vessels and a repeated photothrombosis paradigm was subsequently delivered to the ROI. The 1070 nm laser power was set to 100 mW and the 920 nm laser power was set to 20 mW. The blood flow of the target vessel was imaged as acquired images are updated at ∼2 fps. Rapid aggregation of platelets and blood cells was observed in the vessel’s lumen at 51 s (yellow arrow). The clot grew with the aggregation of blood cells, and eventually blocks blood flow (red arrowhead, at 151.5 s). Following occlusion, the upstream segment of the clot site had slight vasodilation due to the poor flow. The line-scan image was then employed to confirm the occlusion (see Methods, Fig. 3e). In the baseline line-scan image, blood cells moved through the scanning line (orange line), producing streaks (shadow lines), and the slope of the streaks was equal to the inverse of the speed. The line scan image of the occluded vessel appeared as vertical streaks. We further applied the same scan pattern to the no RB-labelled vessels to verify the reliability of our approach. Not surprisingly, there was no sign of blood cells aggregation and clot formation in the control vessel. The line scan image showed no significant flow change was evoked by spiral scanning of the 1070 nm laser for 300s (Fig. 3f, Supplementary Movie 4).

To further compare the efficacy of the raster scan and spiral scan at 1070 nm laser in deep cortical regions, we next explored the capillary photothrombosis from 380 to 400 μm of two scan models (Fig. 4a). We applied raster scan (constant raster scan) (Fig. 4b) and spiral scan (2 s spiral scan following 3 s imaging) (Fig. 4c) in 36 capillaries of 6 mice. The laser power was set to 140 mW in both two scan patterns. There is a significant difference between the effeteness of the two scan patterns, and the spiral scan displayed a more reliable and efficient RB-excitation effect. The spiral scan showed a high success rate of 90%, while the raster scan was difficult to produce occlusion at this depth with a low success rate of 56.25% (Fig. 4d). It is worth noting that in the successful attempts, different from the raster scan, the spiral scan indicated the efficiency of RB-excitation, showing a significant shorter occlusion time (Raster scan: 152.7 ± 66.88 s; spiral scan: 68.06 ± 57.42 s; Mean ± SD; p = 0.0001) (Fig. 4e).

**Figure 4.**
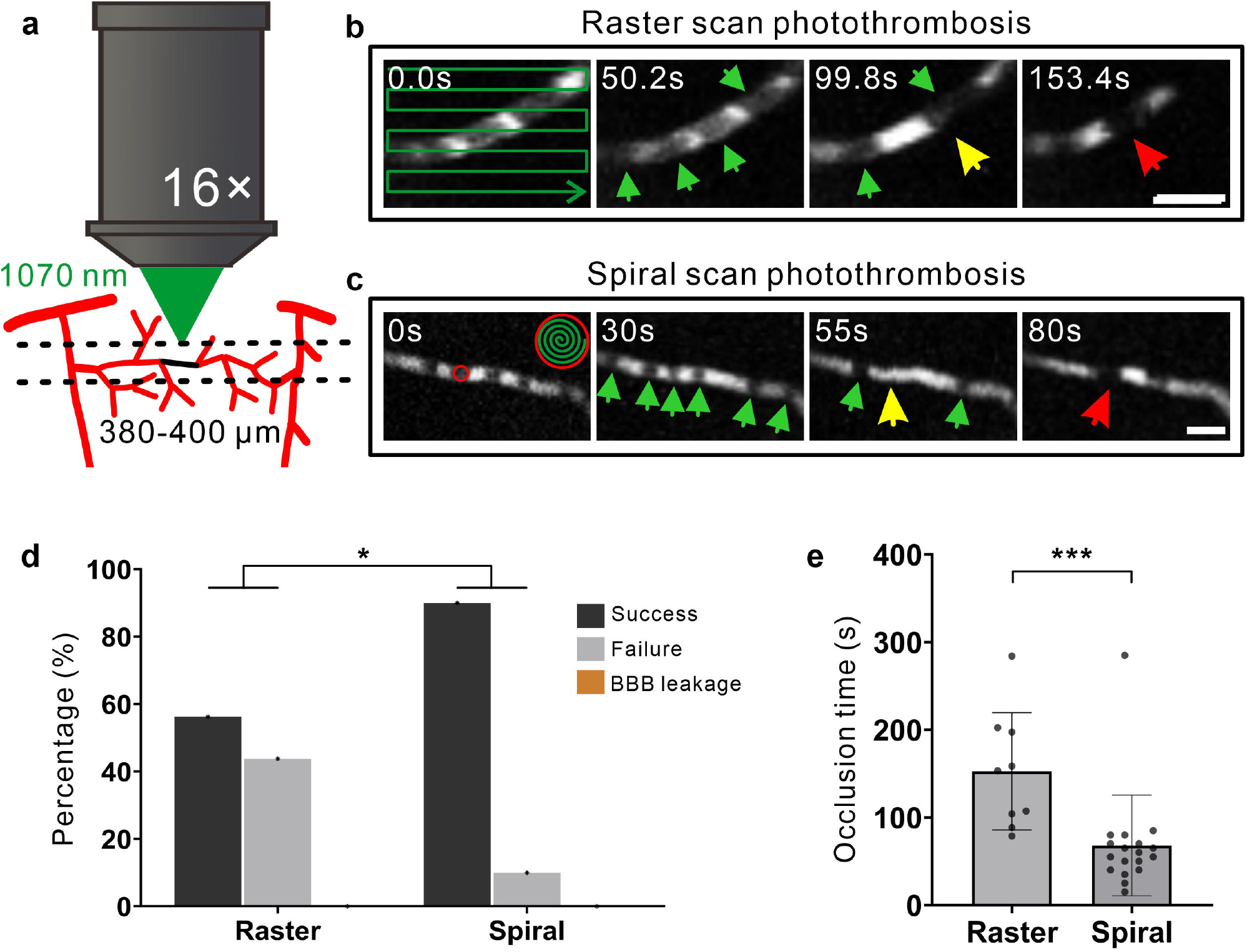
Comparison of the photothrombosis effects between raster and spiral scan (PLP) in 1070 nm. a. Schematic, 1070 nm laser (green) was focused by the objective and irradiated the target vessels in the 380-400 μm cortical layer. b,c. Photothrombosis examples of raster scan (planar b) and spiral scan (planar c) in the target ROI. The green lines in the first slices indicated the scan path. The red circle in the first slice in planar c displayed the target ROI to deliver spiral scans. The green arrows indicated the red blood cells which passed through the vessels’ lumen as black plaques. The yellow arrows indicated the platelets aggregation and attachment to the vessel wall, with a clot (red arrow) formed at 153.4 s and 80 s separately. Scale bar: 10 μm. d. Column plot of percentage of three categories (success, failure, BBB leakage) during capillaries photothrombosis compared between raster and spiral scan. (N= 36 vessels from 5 mice, p = 0.0491, Fisher’s Exact Test) e. Column plot of capillaries occlusion time compared between raster and spiral scan. Each data dot represented the occlusion time of each capillary. (N= 27 vessel from 5 mice, Mean± SD, p= 0.0001, unpaired Mann-Whitney t-test)

### Broad applications of PLP approach for various types of vessels and awake animals

Credit to the great tissue penetration ability of the 1070 nm ultrafast laser and efficiency spiral scan model, our PLP approach provides highly precise and reproducibility photothrombosis over multiple depths and vessel sizes of various types. As the examples shown in Fig. 5a, b, a pial artery and a parenchymal arteriole were occluded via 1070 nm PLP method. The spiral scan was set to apposite power and sizes according to the vessel diameter and depth for photothrombosis (see Methods). The clots appeared in the target segments as the platelet and blood cells aggregation, which can be observed in the planar imaging with TPSLM.

**Figure 5.**
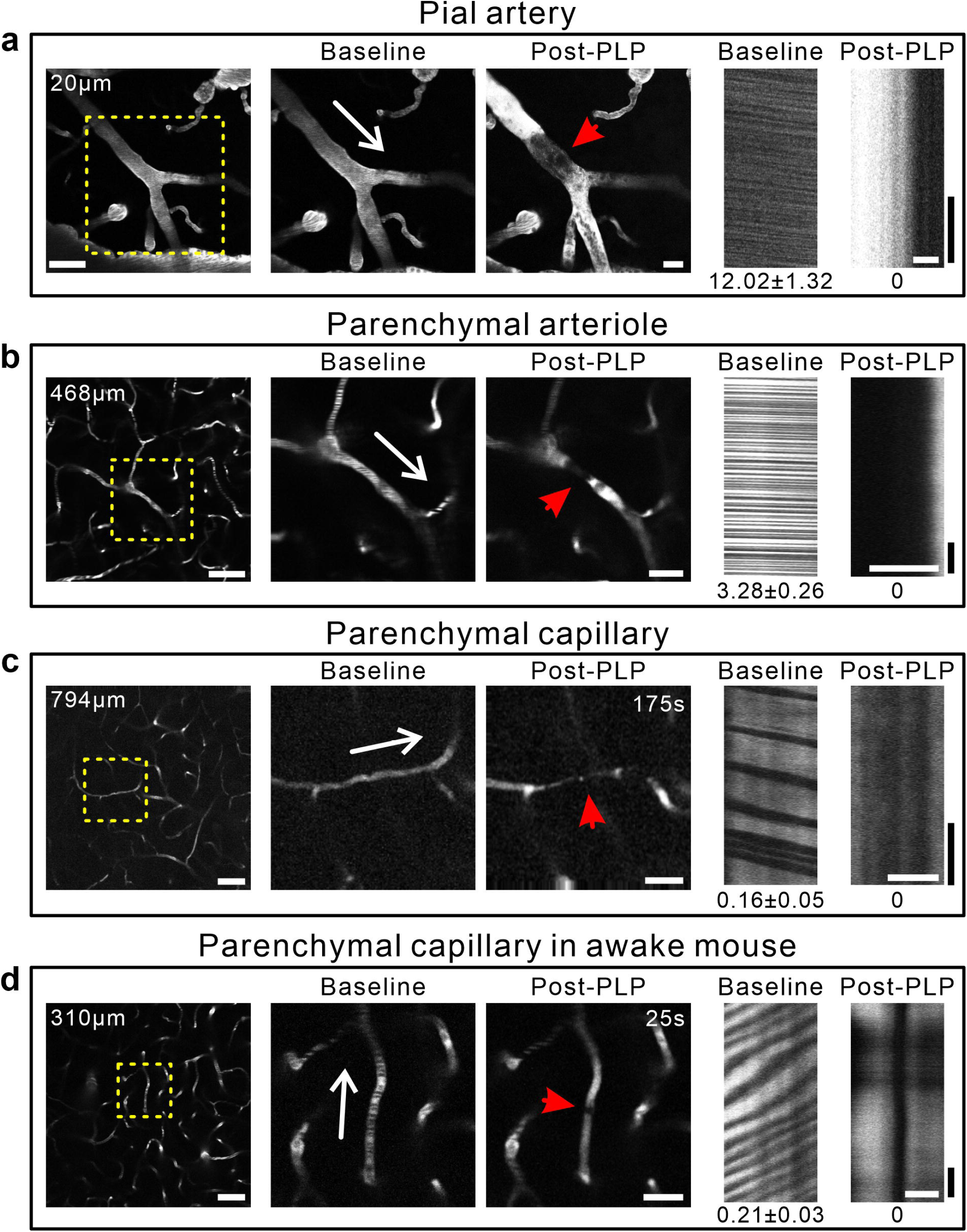
Demonstration of PLP of various vessel types and depths in anesthetized and awake mice. a,b,c. TPLSM images of photothrombotic clotting (red arrow) of a pial artery at 20 μm depth (planar a), parenchymal arteriole at 468 μm depth (planar b), and parenchymal capillary at 794 μm depth (planar c) enlarged from the yellow rectangle in left panel in an anesthetized mouse. d. Parenchymal capillary photothrombosis at 310 μm depth in an awake mouse. The clotted vessels marked with red arrow with bright fluorescence accumulated in the post-thrombosis image. The white arrow corresponds to the flow direction. Line-scan measurements of the flow velocity of the target vessel showed in the right two images. The number below each line-scan image indicated the flow velocity. Unit, mm s^-1^. Scale bar, left: 50 μm; middle: 20 μm; right: horizontal scale bar, 10 μm; vertical scale bar, 0.1s.

Next, we demonstrate a single capillary photothrombosis at a depth of 794 μm (Fig. 5c, Supplementary Movie 5). The laser power was set to 200 mW to induce occlusion at this depth. Accordingly, within 175 s, the blood flow stalled and a clot appeared, which was visible and verified with line-scan imaging. Notably, there were no visible clots in the off-target planes, proving the spatial precision of the RB-excitation (Supplementary Movie 6). And our approach could also induce single-vessel occlusion in awake animals (Fig. 5d, Supplementary Movie 7). The occlusion was formed within merely 25 s, and the stagnant flow was visible and verified with line-scan imaging. Note that there was no discernible extravasation of FITC-dye, indicating the reliability and precision of our method.

We then evaluated the performance of our PLP methods in different vessel types and depths in 20 mice. According to the vessel types and depths, we formulated specific scanning parameters for effective and precision occlusion. We cataloged all vessels into 6 groups according to the differences in vessel morphology types and depths and, found that each group showed diverse “responses” to our method (Fig 6, Table 1). All groups showed high success rates of photothrombosis (Success rate, pial A& DA: 71.43%; parenchymal arteriole: 86.67%; pial V& AV: 92.86%; capillary, 0-200 μm: 93.33%; capillary, 200-500 μm: 95.35%; capillary, below 500 μm: 97.06%), while a few proportions of failure attempts and BBB leakage were observed in all group except below 500 μm group (Failure rate, pial A& DA: 20%; parenchymal arteriole: 6.67%; pial V& AV: 3.57%; capillary, 0-200 μm: 2.22%; capillary, 200-500 μm: 2.33%; capillary, below 500 μm: 2.94%; BBB leakage rate, pial A& DA: 8.57%; parenchymal arteriole: 6.67%; pial V& AV: 3.57%; capillary, 0-200 μm: 4.44%; capillary, 200-500 μm: 2.32%; capillary, below 500 μm: 0%) (Fig. 6a). Notably, the pial artery and diving artery were not easy to form occlusions, as the aggregated platelets and blood cells could be easily rushed away by the turbulent flow. Accordingly, high laser power (100-150 mW) and much more occlusion time were necessary to induce occlusions (Occlusion time, pial A& DA: 174.0 ± 89.37 s; Mean ± SD) (Fig. 6b). Parenchymal arterioles, also pial and ascending veins, which showed much slower blood flow, were much easier to format clots using lower-power energy (80-120 mW) (Occlusion time, parenchymal arteriole: 115.8 ± 84.38 s; pial V& AV: 96.15 ± 45.57 s; Mean ± SD) (Fig. 6b). As capillaries are widely distributed throughout the cortical depth, precise and specific paradigm was customized to adapt the depths demand. Considering the energy loss while laser penetrating the tissue, we set three laser power sections crossing the depths: (1) 0-200 μm: 80-100 mW; (2) 200-500 μm: 100-150 mW; (3) below 500 μm: 150-200 mW. As a result, capillaries showed reliable and fast photothromsbosis, while occlusion time increased with depth (Occlusion time, 0-200 μm: 57.07 ± 32.36 s; 200-500 μm: 88.60 ± 62.33 s; below 500 μm: 111.4 ± 61.79; Mean ± SD) (Fig. 6b). Thus, our PLP method is capable for consistent and effective photothrombosis targeting various types of vessels in a spatially and temporally precise manner.

**Table 1.** PLP parameters for vascular of various sizes and depth.

| Vessel Type | Laser Power (mW) | Revolution | Total Time (s) | Success Rate | 2h maintain |
| --- | --- | --- | --- | --- | --- |
| pial A& DA | 120-150 | spiral<br>0.5μm/circle | 174±89.37 | 71.43% ( 25 of 35 ) | 16 of 25 |
| pial V& AV | 100-120 | | 96.15±45.57 | 92.86% ( 26 of 28 ) | 21 of 26 |
| parenchymal arteriole | 100-120 | | 115.8±84.39 | 86.67% (13 of 15) | 12 of 13 |
| capillary (0-200μm depth) | 80-100 | | 57.07±32.36 | 93.33% (42 of 45) | 41 of 42 |
| capillary (200-500μm depth) | 100-150 | | 88.60±62.33 | 95.35% (41 of 43) | 41 of 41 |
| capillary (below 500μm depth) | 150-200 | | 111.4±61.79 | 97.06% (33 of 34) | 33 of 33 |

**Figure 6.**
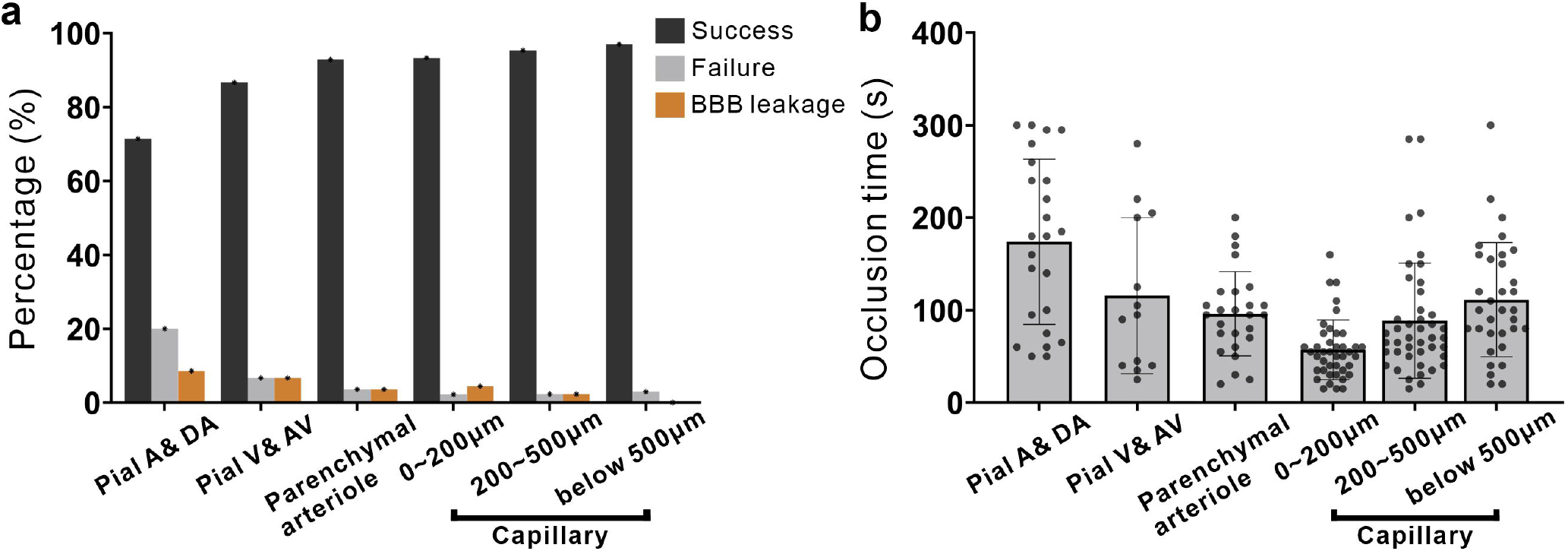
Quantification of the success rate and occlusion time of PLP between different vessel types and depths. a. Column plot of percentage of three categories (success, failure, BBB leakage) vessels after up-to-300 s PLP compared between different vessel types and depths. (N= 200 vessels from 20 mice) b. Column plot of occlusion time compared between different vessel types and depths. Each data dot represented the occlusion time of an individual vessel. (N= 180 vessels from 20 mice, Mean± SD.)

## Discussion

In this study, we demonstrate that RB-mediated photothrombosis is feasible *in vivo* upon excitation of ultrafast lasers at multiple wavelengths. Within 300-s focused raster scan is sufficient for generating microvessel occlusions at a success rate >80% for capillaries located in depths of 80 to 100□μm. Based on this, we established a highly flexible and reproducible PLP method with excitation of spiral scan with 1070 nm ultrafast laser. Compared with conventional approaches, it allows faster, more precise and well-controlled occlusion in various types and depths of vessels with a high success rate while imaging in real time. Non-linear absorption-based photothrombosis prevented damage to the vicinity of neighbor and overlying vessels, promising to mimetic the single vessel dysfunction. Our approach also could induce microvessel occlusions in awake animals, which allows eliminating confounding factors associated with anesthetics. In sum, our approach provides a practical, precise, and depth-selected single-vessel photothrombosis technology with commercially available optical equipment, which is critical for understanding vasculature function in health and disorders.

Traditional RB photothrombosis is typically excited with a green light of 530 to 580 nm to produce an occlusion due to its one-photon absorption peak at around 560 nm^27,28^. Until recently, attention has been attracted that RB can be excited by long-wavelength ultrafast lasers, permitting the approach of precision parenchymal photothrombosis *in vivo*^31,32^. These methods demonstrate the RB-induced photothrombosis with 720 nm or 1000 nm ultrafast laser excitation. Nevertheless, whether the ROS production of RB correlates to the fluorescence emission, and the effect of different wavelengths of ultrafast laser has not been explored. In the current work, we proved that ultrafast light at 720, 780, 860, 920, 1000, and 1070 nm which are widely utilized in TPLSM imaging, are highly efficient and reliable to induce RB photoactivation and subsequent occlusion with a high success rate of above 80% in superficial layer with 80 mW laser power. However, in deep 380-400 μm layer, lasers with 140 mW did not induce efficient occlusion formation due to energy decay upon tissue penetration (success rate below 60%). Notably, we observed fluorescence signals of RB with excitation of all wavelengths of lasers, however, the efficiency of RB-excitation and the resulting occlusion times are different between laser wavelengths. Although fluorescence emission and ROS generation were not systemically quantified in this study, the feasibility of ultrafast laser-induced RB-excitation and subsequent occlusion is encouraging for such a study.

When the laser penetrates through the cortical parenchyma, its energy decays rapidly due to the tissue scattering and water absorption^38^. High-power lasers are critical for RB excitation and subsequent occlusion within deep cortical vessels, however, it would lead to more out-of-focus excitation accompanied by undesired photothrombosis and photothermic damage. In this study, the optimum wavelength window in terms of tissue penetration is 1070 nm when both tissue scattering and absorption are considered. When comparing the photothrombosis effeteness of different wavelengths, 1070 nm laser showed efficiency and reliability, as it showed a fast occlusion formation period and obtained a high success rate of 87.5/56.25% during raster scan excitation.

As the short dwell time and small excitation volume are used in conventional multi-photon scanning, heating through linear absorption can be considered a negligible source of photodamage^40,41^. However, for repetitive raster scanning of restricted ROIs, it draws increasing attention, as it can induce possible photodamages, such as thermal damage related to the linear absorption, multiphoton absorption-induced ablation damage, or optical breakdown^30,42,43^. In the current work, the multi-photon laser was restricted in small ROIs and repeatedly scan at 40-80fps, which may cause photodamage and resulting BBB leakage. Besides this, the scanning in the extraluminal area of the vessel leads to a “discontinuous” RB excitation in the vessel’s lumen, causing an energy “waste” of laser scanning. Although it can be improved by confining the ROIs to the vessels’ lumen, it is unfriendly for real-time imaging to observe clots formation. We employed spiral scan instead of raster scan in the subsequent experiments, which scanned the ROI only in the target vessel’s lumen. Although spiral scan enables high efficient heat dissipation, constant scanning on the ROI may lead to inevitable heat build-up^42^. We thus formulated the photothrombosis paradigm of 3-s spiral scanning combined with alternating 2-s imaging, permitting efficient RB excitation while monitoring the clot formation in real time. Therefore, this paradigm permits precise occlusion in the deep cortical region, the depth of which depends on whether the blood flow can be observed and measured in the focused image plane theoretically.

Cerebral Small Vessel Disease (CSVD) is a common cause of progressive disability and cognitive decline in neurovascular disorders^44,45^. We also used this PLP methods to provide a unique means to mimic vessel dysfunction and model micro-ischemia of these small vascular diseases to neurodegeneration *in vivo*^46^. We can manipulate various types, depths, and sizes of vascular occlusion to obtain different infarct pathologies, which facilitates a microinfarct strategy to study brain injury and repair in neurovascular research coupled with live-imaging technology. It represents a precision medicine approach for vascular disease treatment on a per-vessel/per-lesion basis. Combined with multi-photon imaging, it holds particular promise for studying vascular diseases in the brain cortex, where high spatial selectivity is critical for preventing collateral effects on central nervous system function.

## Methods

### Animal and surgery

All experimental procedures were approved by the Institutional Animal Care and Use Committee of Zhejiang University and in accordance with the Institutes of Health Guide for the Care and Use of Laboratory Animals. Male C57BL/6J mice (9-10 weeks old) were housed at 24°C with a normal 12 h light/dark cycle, and fed with water and food.

Mice were pre-anesthetized with isoflurane (5% inhalation, mixed with fresh air, 0.5L/min) and placed in a stereotactic frame while isoflurane was maintained at 1%–2% throughout the surgical procedure. Body temperature was maintained at 37°C with a heating pad. The craniotomy procedure was performed as previously protocol described^47^. A dental drill exposed a craniotomy (∼4 mm in diameter, the center was at 0.3mm anterior, 2.3mm lateral to bregma) in the right sensory cortex. The dura was carefully moved and artificial cerebrospinal fluid (ACSF) was filled the exposed cortex. Lastly, the round coverslip (diameter: 6 mm, and thickness: 0.17 mm) was used to cover the brain tissue, and medical glue was used to seal its edge. Dental cement was subsequently used to fortify a custom-made stainless-steel headpost with screw holes, allowing the mice to be head-fixed while imaged. Mice received a subcutaneous injection of buprenorphine (0.05 mg kg-1) for at least three days post-surgery and were allowed to recover for at least two weeks before they were imaged.

### Two-photon imaging

All animals with cranial windows were mounted and immobilized to a stationary apparatus with a heating pad and were imaged with TPLSM in anesthetized (1-3 % isoflurane) or awake state. 20 minutes before RB injection, mice were retro-orbitally injected with 50 μL FITC-dextran (10 mg mL^-1^, 2 MDa, FD2000S, Sigma) solution to provide a fluorescent angiogram *in vivo*. A customized TPLSM (2P plus, Bruker Corporation) coupled with a tunable femtosecond laser (80 MHZ, 140 fs, Chameleon Ultra II, Coherent Inc. model-locked Ti: Sapphire laser) was used to record the vascular signals. The femtosecond laser generated two-photon excitation laser (720-1080 nm), whose power was controlled by Pockels Cell (EO-PC, Thorlabs Corporation). Dual-channel fluorescent signals (green: 500-550 nm filter, red: 570-620 nm filter) were collected in photomultiplier tubes (PMTs), which traveled through a 16x (numerical aperture = 0.8) water-immersion objective (3 mm WD, N16XLWD-PF, Nikon) from the samples.

While imaging the vasculature and hemodynamics, the wavelength of the imaging laser was kept at 920 nm with low power (10-50mW, under objective). For planar vasculature imaging, the field of view (FOV) of the individual frame was 375.47× 375.47 μm (pixel: 1024× 1024) with a ∼0.5Hz frame rate. For volume structural imaging, the FOV was set to the same size (pixel: 512× 512) with a ∼2 Hz frame rate (z-axis step: 2 μm), in which each frame was repeated four times, and the averaged frame was taken as the structural results. For flow velocity measurement in real-time, line scan imaging was used, in which repetitive line scanned along the axis of the vessel center, at a rate of at least 4000 Hz. The line-scan imaging formed a space-time image in which moving RBCs produced streaks with a slope that is equal to the inverse of the speed. And we typically acquired line scans for at least 10s totally for one vessel and reported the average speed over this period, which helped to eliminate the influence of blood flow changes in resting state under anesthesia.

### Ultra-fast laser-induced photothrombosis

Once the data of vascular structure and flow velocity in baseline were collected, one dose of ∼50 μL RB solution (1.25%, w/v, in sterile saline) was retro-orbitally injected. As RB is cleared from circulation *in vivo* within 5 minutes^35^, we typically performed photothrombosis within 5 minutes after a one-dose injection. Laser power was measured at the exit of the objective using a laser power meter (PM100D, Thorlabs) before scanning.

Raster scan model. When comparing the photothrombosis effects of different wavelengths of lasers, a targeted rectangle ROI merely including the vessel segment’s lumen was drawn under 920 nm excitation with high digital magnification during the baseline imaging period. The excitation light wavelength was set to a subject wavelength (from 720 nm to 1070 nm), and then the focus planar was quickly readjusted to the target vessel with low power (20 mW). The ROI was subsequently received repeated raster scans at 40-80 fps with the laser power at 80 mW in all attempts, while imaged in real-time. When a clot appeared in the vessel’s lumen and no blood cells passed through over 10s, the scanning process was stopped.

Spiral scan model. When conducting our PLP approach, the 1070 nm laser (70 MHZ, 55 fs, Fidelity 2, Coherent) was directed into the microscope through an individual galvanometer. The laser power was regulated by the Pockels Cells (Conoptics) and was controlled from 80mW to 200mW altering to the specific situation. We formulated a repeated PLP paradigm of 3s scanning followed by 2s imaging to deliver the spiral scan to the target ROI while imaging in real-time. The circled ROI was drawn inside the target vessel’s lumen and was tangent to the vessel wall. When marking spiral scans, the laser starts at the center and, traveling at a constant ultra-fast speed throughout the chosen duration, spirals outward to the edge circling about the center for the number of revolutions specified^30^. The size of the spiral (same as the diameter of the targeted vessel), the number of revolutions (how many revolutions were made to make it from the center of the spiral to the edge, usually interval 0.5 μm per circle), scanning duration (3 s in our paradigm) and repetition (up to 60, 300 s) were customized for different subject vessels. For different vessel types and depths, we have formulated corresponding laser powers to produce reliable and efficient occlusion (Table 1). During the imaging periods of PLP duration, laser power can be adjusted according to our parameters range, which will effectively help to improve the success rate. In addition, the FOV of image can be adjusted according to the experimental demands, which was usually set to 281.61 * 281.61 μm in our experiments. Once the clot was formatted over 10 s, the scanning process stopped subsequently, and line scan was employed to confirm the occlusion.

### Vessels classification

When capillary responses were examined on a vessel-by-vessel basis over the post-thrombosis imaging period, we divided the vessels into three types: (1) success, clot formation and blood flow stalled; (2) failure, blood flow kept constant flow or recovered from stall; (3) BBB leakage, extravasation of FITC or RB around the target blood vessels was discernible during photothrombosis processing.

### Image analysis

All data analyses were performed using MATLAB (R2020b; MathWorks) and Fiji ImageJ (ImageJ, US). Blood flow velocity measurement was carried out by LS-PIV in MATLAB. The slope was calculated using an automated image-processing algorithm^48^. For diameter measurements, each vascular segment was measured manually across the lumen at the peak intensity of the profile using Vasometrics^49^ in Fiji ImageJ software, which was calculated as the FWHM of the intensity profile across the capillary width.

### Statistical Analysis

All statistical analyses were performed using GraphPad Prism (GraphPad Software, San Diego, CA, USA) software. Success and failure attempts comparison between two layers with different wavelengths in Fig. 2e and between two scan patterns in Fig. 4d was analyzed with Chi-Square Fisher’s Exact Test. Occlusion time between raster and spiral scan in Fig. 4b was analyzed with unpaired Mann-Whitney t-test. Results were expressed as mean□± standard deviation, SD. P<□0.05 was accepted as statistically significant. *P< 0.05, **P< 0.01, ***P< 0.001 and ****P< 0.0001.

## Acknowledgements

This project was financially supported by:

National Key R&D program of China 2018YFA0701400 (A.W.R)

National Natural Science Foundation of China 91632105 (W.X.)

National Natural Science Foundation of China U20A20221, 81961128029, 31627802 (A.W.R)

Key R&D Program of Zhejiang 2022C03096 (W.X.)

Key Research and Development Program of Zhejiang Province 2020C03004 (A.W.R)

Fundamental Research Funds for the Central Universities 226-2022-00083 (W.X.)

Fundamental Research Funds for the Central Universities 2019XZZX003-20 (A.W.R).

## Author Contributions

Conceptualization: A.W.R., W.X., L.Z.

Methodology: L.Z., W.X., A.W.R

Investigation: L.Z., M.W., W.X.

Visualization – figures: L.Z., W.X., A.W.R., M.W., P.F., Y. L., H.Z.

Funding acquisition: W.X., A.W.R.

Project administration: W.X., A.W.R., L.Z.

Supervision: W.X., A.W.R.

Writing – original draft: L.Z., W.X., A.W.R.

Writing – review & editing: L.Z., W.X., A.W.R., M.W., Y.L., P.F.

Data Analysis: L.Z., W.X.

## Declaration of interests

The authors declare no competing interests.

## Notes

### Competing Interest Statement

The authors have declared no competing interest.

## References

1. Iadecola, C. The Neurovascular Unit Coming of Age: A Journey through Neurovascular Coupling in Health and Disease. Neuron 96, 17–42 (2017).

2. Iadecola, C. The pathobiology of vascular dementia. Neuron 80, 844–866 (2013).

3. Girouard, H. & Iadecola, C. Neurovascular coupling in the normal brain and in hypertension, stroke, and Alzheimer disease. J Appl Physiol (1985) 100, 328–335 (2006).

4. Blinder, P. et al. The cortical angiome: an interconnected vascular network with noncolumnar patterns of blood flow. Nat Neurosci 16, 889–897 (2013).

5. Kirst, C. et al. Mapping the Fine-Scale Organization and Plasticity of the Brain Vasculature. Cell 180, 780–795 e725 (2020).

6. Ji, X. et al. Brain microvasculature has a common topology with local differences in geometry that match metabolic load. Neuron 109, 1168–1187 e1113 (2021).

7. Chen, B. R., Bouchard, M. B., McCaslin, A. F., Burgess, S. A. & Hillman, E. M. High-speed vascular dynamics of the hemodynamic response. Neuroimage 54 (2011).

8. Ances, B. M. Coupling of changes in cerebral blood flow with neural activity: what must initially dip must come back up. J Cereb Blood Flow Metab 24, 1–6 (2004).

9. Hillman, E. M. Coupling mechanism and significance of the BOLD signal: a status report. Annu Rev Neurosci 37, 161–181 (2014).

10. Uludag, K. & Blinder, P. Linking brain vascular physiology to hemodynamic response in ultra-high field MRI. Neuroimage 168, 279–295 (2018).

11. Grinvald, A., Lieke, E., Frostig, R. D., Gilbert, C. D. & Wiesel, T. N. Functional architecture of cortex revealed by optical imaging of intrinsic signals. Nature 324, 361–364 (1986).

12. Lu, H. D., Chen, G., Cai, J. & Roe, A. W. Intrinsic signal optical imaging of visual brain activity: Tracking of fast cortical dynamics. Neuroimage 148, 160–168 (2017).

13. Sweeney, M. D., Kisler, K., Montagne, A., Toga, A. W. & Zlokovic, B. V. The role of brain vasculature in neurodegenerative disorders. Nat Neurosci 21, 1318–1331 (2018).

14. Cruz Hernandez, J. C. et al. Neutrophil adhesion in brain capillaries reduces cortical blood flow and impairs memory function in Alzheimer’s disease mouse models. Nat Neurosci 22, 413–420 (2019).

15. Greenberg, S. M. et al. Cerebral amyloid angiopathy and Alzheimer disease - one peptide, two pathways. Nat Rev Neurol 16, 30–42 (2020).

16. Zlokovic, B. V. The blood-brain barrier in health and chronic neurodegenerative disorders. Neuron 57, 178–201 (2008).

17. Logothetis, N. K., Pauls, J., Augath, M., Trinath, T. & Oeltermann, A. Neurophysiological investigation of the basis of the fMRI signal. Nature 412, 150–157 (2001).

18. Sirotin, Y. B. & Das, A. Anticipatory haemodynamic signals in sensory cortex not predicted by local neuronal activity. Nature 457, 475–479 (2009).

19. Durukan, A. & Tatlisumak, T. Acute ischemic stroke: Overview of major experimental rodent models, pathophysiology, and therapy of focal cerebral ischemia. Pharmacol Biochem Be 87, 179–197 (2007).

20. Longa, E. Z., Weinstein, P. R., Carlson, S. & Cummins, R. Reversible Middle Cerebral-Artery Occlusion without Craniectomy in Rats. Stroke 20, 84–91 (1989).

21. Luo, W. H., Wang, Z., Li, P. C., Zeng, S. Q. & Luo, Q. M. A modified mini-stroke model with region-directed reperfusion in rat cortex. J Cerebr Blood F Met 28, 973–983 (2008).

22. Silasi, G. & Murphy, T. H. Stroke and the connectome: how connectivity guides therapeutic intervention. Neuron 83, 1354–1368 (2014).

23. Jia, J. M. et al. Control of cerebral ischemia with magnetic nanoparticles. Nat Methods 14, 160–166 (2017).

24. Schmidt, R., Schmidt, H. & Fazekas, F. Vascular risk factors in dementia. Journal of Neurology 247, 81–87 (2000).

25. Nishimura, N. et al. Targeted insult to subsurface cortical blood vessels using ultrashort laser pulses: three models of stroke. Nat Methods 3, 99–108 (2006).

26. Huang, Y. et al. Precise closure of single blood vessels via multiphoton absorption-based photothermolysis. Sci Adv 5, eaan9388 (2019).

27. Zhang, S., Boyd, J., Delaney, K. & Murphy, T. H. Rapid reversible changes in dendritic spine structure in vivo gated by the degree of ischemia. J Neurosci 25, 5333–5338 (2005).

28. Watson, B. D., Dietrich, W. D., Busto, R., Wachtel, M. S. & Ginsberg, M. D. Induction of reproducible brain infarction by photochemically initiated thrombosis. Ann Neurol 17, 497–504 (1985).

29. Watson, B. D., Prado, R., Veloso, A., Brunschwig, J. P. & Dietrich, W. D. Cerebral blood flow restoration and reperfusion injury after ultraviolet laser-facilitated middle cerebral artery recanalization in rat thrombotic stroke. Stroke 33, 428–434 (2002).

30. Picot, A. et al. Temperature Rise under Two-Photon Optogenetic Brain Stimulation. Cell Rep 24, 1243–1253 e1245 (2018).

31. Delafontaine-Martel, P. et al. in Optical Techniques in Neurosurgery, Neurophotonics, and Optogenetics (2021).

32. Fukuda, M., Matsumura, T., Suda, T. & Hirase, H. Depth-targeted intracortical microstroke by two-photon photothrombosis in rodent brain. Neurophotonics 9, 021910 (2022).

33. Machler, P. et al. In Vivo Evidence for a Lactate Gradient from Astrocytes to Neurons. Cell Metab 23, 94–102 (2016).

34. Roche, M. et al. In vivo imaging with a water immersion objective affects brain temperature, blood flow and oxygenation. Elife 8 (2019).

35. Zhang, S. & Murphy, T. H. Imaging the impact of cortical microcirculation on synaptic structure and sensory-evoked hemodynamic responses in vivo. PLoS Biol 5, e119–e119 (2007).

36. Danielczok, J. G. et al. Red Blood Cell Passage of Small Capillaries Is Associated with Transient Ca(2+)-mediated Adaptations. Front Physiol 8, 979 (2017).

37. Wu, X. et al. Tether-free photothermal deep-brain stimulation in freely behaving mice via wide-field illumination in the near-infrared-II window. Nat Biomed Eng (2022).

38. Horton, N. G. et al. In vivo three-photon microscopy of subcortical structures within an intact mouse brain. Nat Photonics 7 (2013).

39. Bashkatov, A. N., Genina, E. A., Kochubey, V. I. & Tuchin, V. V. Optical properties of human skin, subcutaneous and mucous tissues in the wavelength range from 400 to 2000□nm. Journal of Physics D: Applied Physics 38, 2543–2555 (2005).

40. Debarre, D., Olivier, N., Supatto, W. & Beaurepaire, E. Mitigating phototoxicity during multiphoton microscopy of live Drosophila embryos in the 1.0-1.2 microm wavelength range. PLoS One 9, e104250 (2014).

41. Linz, N., Freidank, S., Liang, X.-X. & Vogel, A. Wavelength dependence of femtosecond laser-induced breakdown in water and implications for laser surgery. Physical Review B 94 (2016).

42. Hopt, A. & Neher, E. Highly Nonlinear Photodamage in Two-Photon Fluorescence Microscopy. Biophysical Journal 80, 2029–2036 (2001).

43. Vogel, A., Noack, J., Hüttman, G. & Paltauf, G. Mechanisms of femtosecond laser nanosurgery of cells and tissues. Applied Physics B 81, 1015–1047 (2005).

44. Pantoni, L. Cerebral small vessel disease: from pathogenesis and clinical characteristics to therapeutic challenges. Lancet Neurol 9, 689–701 (2010).

45. Wardlaw, J. M., Smith, C. & Dichgans, M. Mechanisms of sporadic cerebral small vessel disease: insights from neuroimaging. Lancet Neurol 12, 483–497 (2013).

46. Zhu, L. et al. Imaging microvasculature network evolution and neurodegeneration with precise photothrombosis approach. bioRxiv, 2021.2011.2029.470313 (2021).

47. Goldey, G. J. et al. Removable cranial windows for long-term imaging in awake mice. Nat Protoc 9, 2515–2538 (2014).

48. Kim, T. N. et al. Line-scanning particle image velocimetry: an optical approach for quantifying a wide range of blood flow speeds in live animals. PLoS One 7, e38590 (2012).

49. McDowell, K. P., Berthiaume, A. A., Tieu, T., Hartmann, D. A. & Shih, A. Y. VasoMetrics: unbiased spatiotemporal analysis of microvascular diameter in multi-photon imaging applications. Quant Imaging Med Surg 11, 969–982 (2021).

